# 2′-5′-Oligoadenylate synthetase-like protein inhibits intracellular *M. tuberculosis* replication and promotes proinflammatory cytokine secretion

**DOI:** 10.1101/821173

**Authors:** G. Leisching, A. Ali, V. Cole, B. Baker

## Abstract

Host cytoplasmic surveillance pathways are known to elicit type I interferon (IFN) responses which are crucial to antimicrobial defense mechanisms. Oligoadenylate synthetase-like (OASL) protein has been extensively characterized as a part of the anti-viral mechanism, however a number of transcriptomic studies reveal its upregulation in response to infection with a wide variety of intracellular bacterial pathogens. To date, there is no evidence documenting the role (if any) of OASL during mycobacterium tuberculosis infection. Using two pathogenic strains differing in virulence only, as well as the non-pathogenic *M. bovis* BCG strain, we observed that pathogenicity and virulence strongly induced *OASL* expression after 24 h of infection. Further, we observed that *OASL* knock down led to a significant increase in *M. tb* CFU counts 96 h post-infection in comparison to the respective controls. Luminex revealed that OASL silencing significantly decreased IL-1β, TNF-α and MCP-1 secretion in THP-1 cells and had no effect on IL-10 secretion. We therefore postulate that OASL regulates pro-inflammatory mediators such as cytokines and chemokines which suppress intracellular mycobacterial growth and survival.

## Introduction

Susceptibility to tuberculosis encompasses a complex myriad of host responses, which when faced with infection by *Mycobacterium tuberculosis* (*M. tb*), ultimately fail to contain and eliminate the pathogen. The clarification of susceptibility to tuberculosis which is underpinned by genetic control, is expected to provide effective tools for disease prevention and eventually, personalised host-directed therapeutic options. To date however, many of the genes and mechanisms that determine susceptibility to tuberculosis still remain unidentified.

Intracellular host defence mechanisms in particular play a fundamental role in the initial innate immune response and containment of *M. tb* infection in the macrophage, whether it be viral or bacterial. Oligoadenylate synthetise-like (OASL) protein has been extensively characterised as a part of the host defence mechanism against viral infections. *OASL* is induced by both type I and type II interferons (IFN) [1], but independently of this, is also a nucleic acid sensor which can be activated upon binding with foreign or self dsRNA or dsDNA[2, 3]. In response to infection with RNA viruses, OASL functions as an anti-viral protein through enhancing signalling through the RIG-1 pathway and stimulating IFNαβ production [4]. However during infection with DNA viruses, OASL inhibits the cyclic GMP-AMP (cGAMP) synthase (cGAS) sensor which consequently limits the type I IFN response [2]. The role of OASL therefore bifunctional and demonstrates both an IFN inhibitory and promoting function within the cell which is indicative of its ability to differentially modulate Type I IFN production during DNA and RNA viral infection [5].

The question that arises is whether this protein may function in a similar manner when sensing foreign dsDNA from intracellular pathogens such as mycobacteria. Evidence from a number of transcriptomic studies reveal that *OASL* is upregulated in various host cell types infected with a wide range of intracellular gram-positive, gram-negative and acid-fast bacterial pathogens as early as 4 hour post-infection [6-12]. Whether it has as a pro- or anti-bacterial function was unconfirmed until de Toledo and colleagues observed that OASL promoted *M. leprae* intracellular survival, which they suggest is associated with the ability of OASL to inhibit autophagic mechanisms which prevented the clearance of the bacteria [13]. To date, there is no evidence documenting the role (if any) of OASL during *M. tuberculosis* infection.

To address this, we firstly investigated and characterised the expression and biological role of OASL in response to pathogenic and non-pathogenic mycobacterial infection in THP-1 macrophages. The objectives were therefore to i) characterise whether the expression of *OASL* is specific to pathogenicity and/or virulence of mycobacteria, ii) determine whether OASL knock-down affects intracellular survival of mycobacteria, and ii) determine whether OASL expression affects cytokine secretion during the infection state.

## Materials and Methods

### Cells and culture medium

Human macrophage-like cell line THP-1(ATCC-88081201), were cultured in RPMI-1640 supplemented with 10% heat-inactivated foetal calf serum (Biochrome, Germany) and incubated at 37°C, 5% CO2. Before infection experiments, THP-1s were differentiated into macrophage-like cells with Phorbol 12-Myristate 13-Acetate (PMA; Sigma Aldrich, USA) at a final concentration of 100 nM for 24 hours prior to infection.

### Bacterial strains and infection conditions

Hyper (R5527) and hypovirulent (R1507) Beijing *M. tuberculosis* clinical isolates [14] were used for infection, as well as the non-pathogenic *M. bovis* BCG strain. Mycobacteria were cultured in 7H9 (supplemented with 10% OADC, 0.5% glycerol) without Tween 80 [15, 16]. THP-1s were infected with either the hyper-, hypovirulent *M.tb* or *M. bovis* BCG strain at a MOI=1 using the “syringe settle filtrate” (SSF) method [15] and allowed 4 h for uptake (See supplementary Figure S1 for uptake). The cells were then washed 3 times with phosphate buffered saline (PBS) to remove any extracellular mycobacteria, and incubated for a further 20 or 92 hours in complete medium. Uninfected THP-1s served as the control/uninfected samples. MOIs as well as intracellular replication during infection was assessed through CFU counting. Briefly, after each infection time point THP-1s were lysed with 0.1 % sterile Triton X-100 where after the bacteria were serially diluted in 7H9 and plated out on 7h11 agar for enumeration.

### RNA extraction

Total RNA from THP-1 cells were extracted using the RNeasy® Plus Mini Kit (Cat. No. 74134, Qiagen, Limburg, Netherlands) according to the manufacturer’s instructions at 24 and 96h p.i. The ‘gDNA eliminator’ column included in this kit was used to remove genomic DNA in all samples. For each experiment, RNA quality and quantity was assessed using the Agilent 2100 Bioanalyser. Only RNA with a RNA integrity Number (RIN) above 9.0 were used for qPCR experiments.

### Quantitative PCR

For cDNA synthesis, 0.5 µg RNA was converted into cDNA using the Quantitect® Reverse Transcription Kit (Cat. No. 205311, Qiagen, Limburg, Netherlands). To ensure the removal of genomic DNA, ‘gDNA wipe-out buffer’ was added to RNA (included in the kit) prior to the RNA conversion step. qPCR amplification was performed in 96-well plates and run on a LightCycler® 96 system (Roche, Germany). LightCycler® 480 SYBR Green I Master (Cat. No. 04887352001, Roche, Germany) was used to detect *OASL* expression together with a QuantiTect® primer assay (Qiagen, Limburg, Netherlands) for *OASL* (Cat. No. QT00019796) at a reaction volume of 20 µl. *UBC* (Cat. No.QT00234430) and *G6PD* (Cat. No. QT00071596) were used as reference genes. The amplification procedure entailed 45 cycles of 95 °C for 10 s followed by 60 °C for 10s and finally 72 °C for 10s. Gene expression fold-changes were computed for *M. bovis* BCG, hypovirulent *M.tb* infected, hypervirulent *M.tb* infected and uninfected macrophages using calibrated normalised relative quantities using the equation N = N_0_ x 2^Cp^ (LightCycler®96 software, Roche). All qPCRs were done on RNA extracted from three separate experiments. All biological replicates were run in triplicate with a positive control (calibrator) and a non-reverse transcription control in accordance with the MIQE nGuidelines[17].

### Western blotting

After 24 h of infection, protein was extracted using RIPA buffer (150 mM NaCl, 1.0% Triton X-100, 0.5% sodium deoxycholate, 0.1% SDS and 50 mM Tris) containing Complete Protease Inhibitor Cocktail Tablets (Cat. No. 04693116001, Roche Diagnostics, South Africa). Proteins of interest for human were detected with antibodies (Santa Cruz Biotechnology) specific for OASL (sc-98313) and the reference protein GAPDH (sc-32233). Corresponding secondary antibody used was goat anti-rabbit IgG-HRP (sc-2030).

### *OASL* silencing in THP-1 cells

*OASL* was silenced using the FlexiTube siRNA Premix (Qiagen, Cat. # 1027420) for rapid siRNA transfection. Two siRNAs (Cat. No. SI03068933 and Cat. No. SI00055216) with different target sequences were used for silencing. A siRNA negative (non-silencing) control was included (Cat. No. SI03650325). Addition of the siRNA was done after the 4h uptake period according to the manufacturer’s instructions.

### MTT cell viability assay

To evaluate whether silencing of OASL affected host cell viability after each infection period, the 3-(4,5-dimethylthiazol-2-yl)-2,5-diphenyl tetrazolium bromide (MTT) assay was used. It is based on the ability of viable cells with viable mitochondria to reduce MTT into blue formazan pigments and is therefore used to evaluate cytotoxicity [18]. After 24 and 96 h, 0.01 g/ml MTT was added to the cells and incubated for 2 h at 37°C at 5% CO_2_. HCl-isopropanol-Triton solution (1% HCl in isopropanol, 0.1% Triton X-100 in a 50:1 ratio) was then added to the cells in order to release any intracellular formazan produced. The optical density (OD) was determined on a plate reader (EL-800, Micro-Tek instruments) at a wavelength of 540 nm and the values expressed as a percentage of the control (uninfected cells).

### Luminex®

After each infection period, cell culture medium was removed and frozen at −80 °C until analysis. The Milliplex® map human cytokine/chemokine magnetic bead panel kit (Cat.# HCYTOMAG-60K) was used to simultaneously quantify the concentrations of TNF-α, IL-1β, IL-10 and MCP-1 in cell culture supernatants at 24 and 96 h post-infection on the Bioplex 200 (Bio-Rad). The Bioplex Manager 6.1 software was used for data analysis (Bio-Rad). Supernatants were collected from three independent experiments, each with 4 technical replicates which were run in duplicate.

### Immunofluorescence and Analysis

THP-1s were seeded on 14mm glass cover slips and infected with either *Mycobacterium bovis* BCG or the hypo or hypervirulent *M. tb* strains, or remained uninfected as described above. Twenty four hours post infection, the cells were heat-fixed at 95°C for 2 hours, followed by permeabilization with 0.2 % Triton X-100 in PBS for 10 minutes. Cells were immunostained with an OASL (Santa Cruz sc-98313) primary antibody in 3% BSA for 24 hours at 4°C. Cells were then incubated with Texas Red secondary antibody (Sigma Aldrich SAB3700837) in 3% BSA in PBS for 90 minutes and nuclei counter-stained with Hoechst for 10 minutes. Finally, cells were washed in PBS and cover slips were mounted on microscope slides using DAKO fluorescent mounting medium. Images were acquired on a LSM 780 Elyra S1 Confocal laser-scanning microscope, with a 63X, 1,4NA, oil-immersion objective at the Central Analytical Facility (Stellenbosch University, RSA). Hoechst was viewed with a 360 DAPI Filter (Intensity 42%, Gain – 500ms) and the Texas Red Marker with a 572 Texas Red filter (Intensity 100%, Gain – 1000ms).

Immunofluorescence was analyzed quantitatively by measuring the fluorescent intensity at each pixel across the images using histogram analysis in Image J (Windows version; National Institutes of Health) [19]. Background signal was eliminated in the images by using the appropriate thresholding before histogram analysis. The results of the analysis of 12 images acquired in each experimental condition, carried out in triplicate, were then combined to allow quantitative estimates of changes in fluorescence.

### Statistical analysis

Statistical significance was performed with GraphPad Prism software. ANOVA was used for comparisons involving 3 or more groups, with a Bonferroni post-hoc correction applied. All values expressed as means ± SEM with a *p* < 0.05 considered as significant.

## Results

### Pathogenic *M. tb* infection induces OASL expression in THP1 macrophages 24 h post-infection

Since OASL has been extensively characterised as an anti-viral protein, we firstly wanted to determine whether Mycobacterial infection induced its expression. Secondly we sought to determine whether pathogenicity and virulence of mycobacteria affected the extent to which it was expressed (Figure 1). After 24 h of infection with both hyper- and hypovirulent strains, we observed a significant increase in OASL mRNA (Fig. 1A). Relative protein levels of OASL were also significantly increased and displayed the same trend (Fig. 1.B), which was confirmed using immunofluorescence (Fig. 1.C). Interestingly, infection with *M. bovis* BCG did not elicit a transcriptional response in this gene, however a longer infection phase may induce *OASL* expression. We therefore confirmed that pathogenic mycobacteria induce OASL expression and that this expression is independent of virulence.

**Figure 1.**
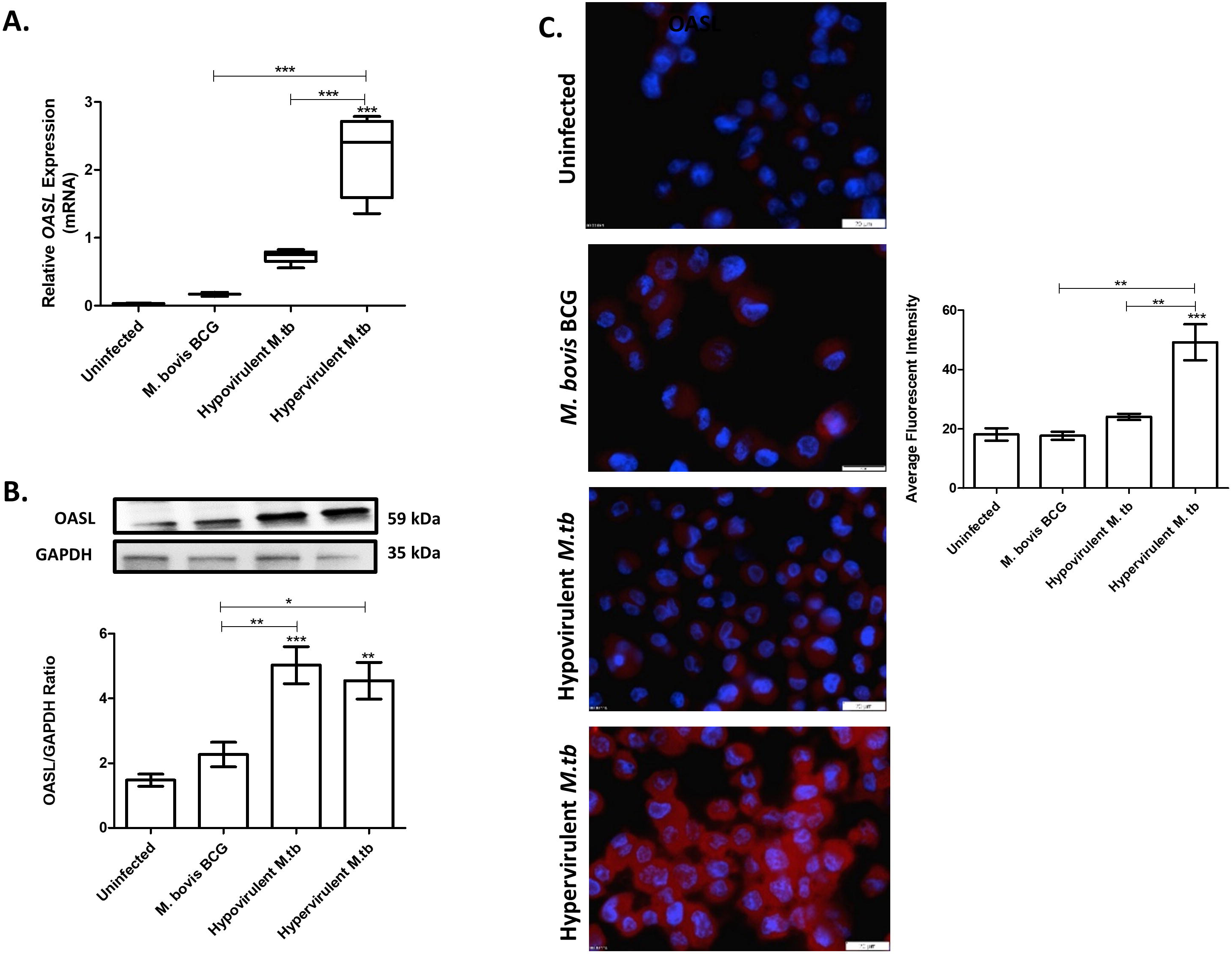
OASL expression in THP-1 cells following infection with M. bovis BCG, R1507 M.tb (hypovirulent) and R5527 M.tb (hypervirulent). A. qPCR analysis of OASL expression 24 h post-infection. UBC and GAPDH were used as reference genes to calculate the relative expression of the target genes. B. OASL expression quantified by Western blotting C. THP-1s were immunostained with OASL primary antibody and then stained with a Texas Red secondary antibody. Nuclei were counter-stained with Hoechst (blue). Average fluorescent intensity was quantified using Image J. 12 images acquired in each experimental condition, carried out in triplicate. Scale bar = 20 μM. One-way ANOVA with a Bonferroni post-hoc test was applied, ** p<0.01, *** p<0.001 vs. uninfected.

### Silencing of *OASL* promotes intracellular replication of pathogenic and non-pathogenic mycobacteria

In order to determine whether the presence of OASL affects the intracellular survival and replication of mycobacteria, we silenced *OASL* using 2 siRNA complexes which each have a different target sequence. Once knock-down was confirmed for both siRNA complexes (Fig. S1.B), THP-1 cells were infected with the three strains and allowed 96 h to establish infection (Figure 2). Host cell viability was also assessed during this time (Fig. 2 A-C). No significant differences in CFUs were observed 24 h post-infection for any of the strains, however after 96 h, intracellular CFU counts increased significantly. *M. bovis* BCG intracellular CFUs increased after *OASL* knock-down with both siRNA target sequences and no significant differences in host-cell viability was observed (Fig. 2A). A similar trend was observed for both hypo- and hypervirulent infection states: siRNA knock-down of *OASL* induced significant increases in intracellular CFUs when compared to the siRNA scrambled sequence (Negative control) and infection without a silencing complex (Fig. 2B and C). These results indicate that OASL exerts a suppressive effect on intracellular mycobacterial replication over time.

**Figure 2.**
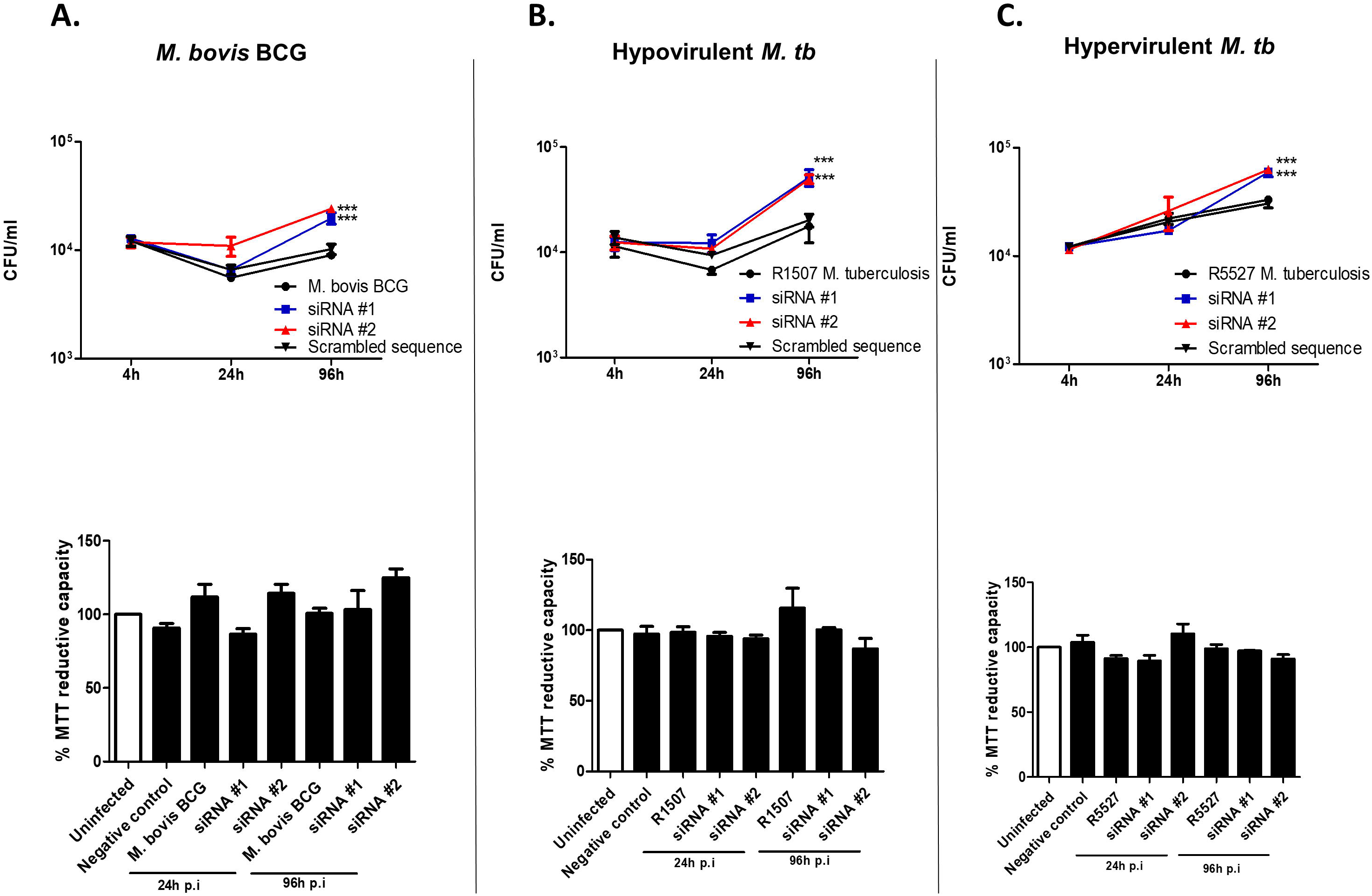
Intracellular growth of pathogenic and non-pathogenic mycobacteria and THP-1 viability after silencing of *OASL* over a 96 h time period. Two siRNA molecules (siRNA #1 and siRNA #2) with different target sequences were used for each gene, as well as scrambled sequence (negative control) **A.** *OASL* silencing did not affect host-cell viability but significantly increased the intracellular CFU counts of *M. bovis* BCG 96 h p.i. **B.** Hypovirulent *M.tb* CFU counts increased significantly 96 p.i. after *OASL* silencing, **C.** Intracellular CFU counts increased significantly 96 h p.i during infection with the hypovirulent *M.tb* strain. No significant effects on THP-1 viability was observed. Two-way ANOVA with a Bonferroni post-hoc test was applied, ** p<0.01, *** p<0.001 vs. infected unsilenced and infected silenced (scrambled sequence), n=4.

### OASL knock-down decreases the secretion of TNFα, IL-1β and MCP-1 from THP-1 cells infected with pathogenic and non-pathogenic mycobacteria

Proinflammatory cytokines act synergistically to control the intracellular replication of mycobacteria within the macrophage [20], and since *OASL* expression is an interferon-induced gene, we sought to determine whether *OASL* silencing affected cytokine/chemokine expression using Luminex® technology. Other cytokines were analysed, however the concentrations after the infection period were too low for detection, and therefore not included in the results (Figure 3).

**Figure 3.**
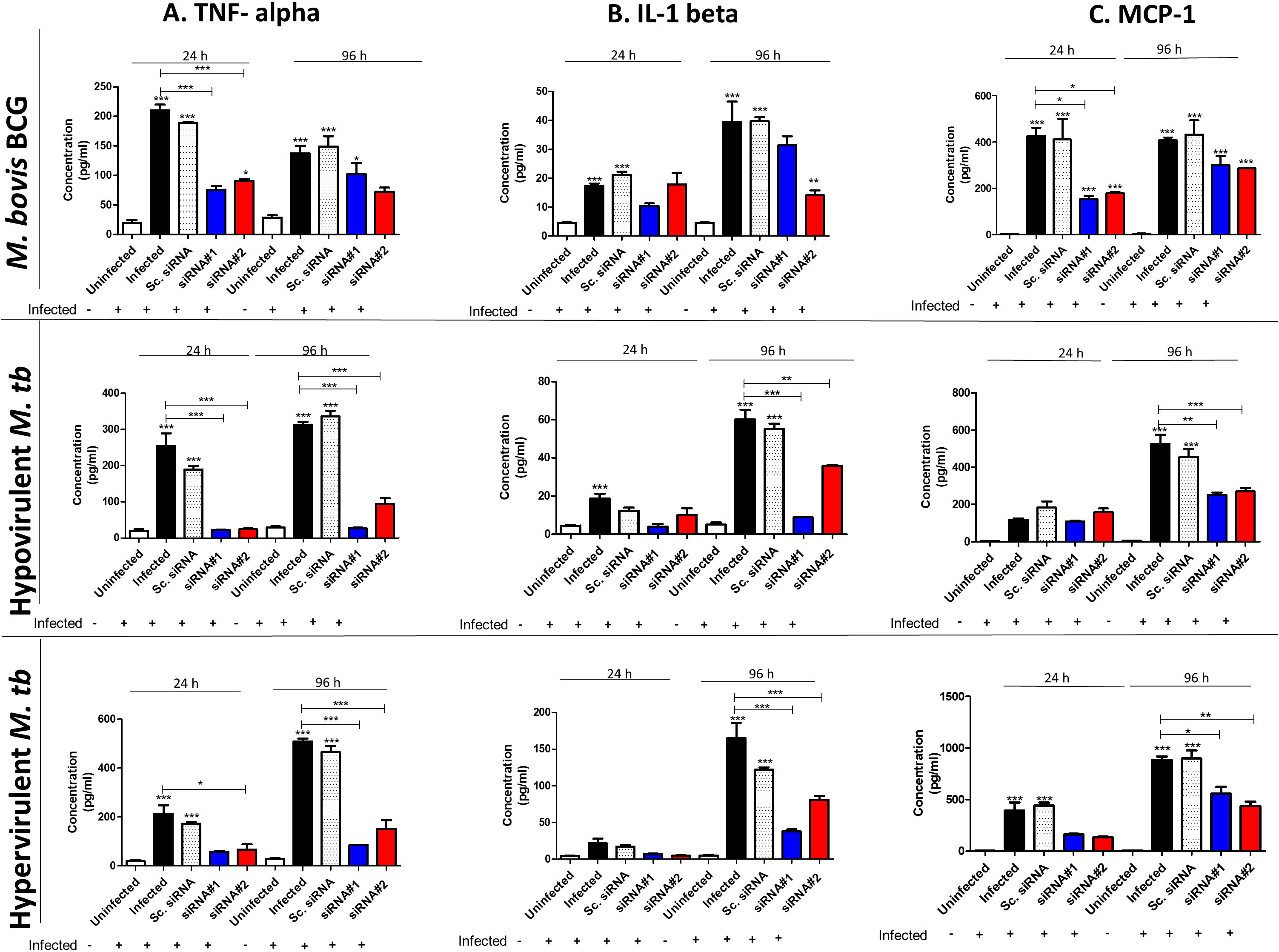
The effect of OASL silencing on TNF-α, IL1-β and MCP-1 secretion in THP-1 cells infected with pathogenic and non-pathogenic mycobacteria as measured by Luminex. Two siRNA molecules (siRNA #1 and siRNA #2) with different targets were used for OASL, as well as scrambled sequence (Sc. siRNA-negative control) A. TNF-α secretion decreased significantly at both 24 and 96 h p.i after OASL silencing in both pathogenic and non-pathogenic infection states. B. Significant differences in IL-1β secretion were only observed 96 h p.i with pathogenic and non-pathogenic mycobacteria. C. MCP-1 decreased significantly 96 h p.i with pathogenic mycobacteria only. One-way ANOVA with a Bonferroni post-hoc test was applied, ** p<0.01, *** p<0.001 vs. uninfected, n=3.

TNFα secretion appeared to be significantly affected by the silencing of *OASL* which was independent of virulence and pathogenicity of the mycobacteria (Fig. 3A). This is an important observation since TNFα plays a prominent role in controlling intracellular mycobacterial replication [21]. TNF-α release was significantly reduced 24 h p.i with both pathogenic and non-pathogenic mycobacterial infection states after OASL silencing in comparison to the unsilenced infected control and the infected scrambled sequence control (Fig.3A). This effect remained after 96 h p.i however only in THP-1s infected with pathogenic mycobacteria. Next we assessed the effects of OASL silencing on IL-1β secretion (Fig.3B). Although infection alone induced significant IL-1β secretion at 24 h, OASL silencing had no effect on this secretion. At 96 h however, IL-1β levels were significantly downregulated in THP-1s infected with both pathogenic strains (Fig. 3C). The analysis of the chemokine MCP-1 (*CCL2*) indicated that only at 24 h p.i with *M. bovis* BCG, was a significant reduction in secretion observed. *OASL* silencing significantly reduced MCP-1 secretion 96 h p.i during infection with pathogenic mycobacteria only. Although we observed detectable levels of IL-10 in our model, they were only detectable 96 h p.i and observed that IL-10 release was unaffected by the silencing of *OASL* (Fig. S1 C).

The effects of *OASL* silencing on cytokine and chemokine secretion in THP-1 macrophages can be summarised as follows: i) TNF-α and IL-1β secretion is enhanced during mycobacterial infection in the presence of *OASL*, ii) OASL enhances MCP-1 production in response to THP-1 infection by pathogenic mycobacteria only, and iii) IL-10 production is unaffected by the presence/absence of *OASL*.

## Discussion

Interferon-inducible oligoadenylate synthetase-like (OASL) protein enhances type I interferon (IFN) induction and inhibits intracellular viral replication. Its expression is as a result of IFNαβ release and thus forms part of the interferon stimulated genes (ISGs) that are associated with the early inflammatory response during infection. It has since been put forward that the anti-viral capability of OASL be harnessed with the potential for developing broad acting antiviral therapy [21]. Interestingly, *OASL* is amongst the set of genes defined as the TB signature that discriminates active from latent tuberculosis and other diseases [22-24] which suggests that this gene may play a role during active mycobacterial infection, although this is yet to be determined. Its upregulation has also been observed in response to a number of Gram-positive and Gram –negative, and acid-fast bacteria from as early as 4 h p.i. These pathogens include *Brucella abortus* [9], *Chlamydia trachomatis* [6], *Lactobacillus acidophilus* [10], *Chlamydia pneumoniae* [7], *Listeria monocytogenes* [8], *Mycobacterium leprae* [13] and in *Mycobacterium tuberculosis* [11] (a previous study in our lab). Questions regarding the biological role of OASL during *M. tuberculosis* infection remained unanswered. This study therefore aimed to characterise the role of OASL in response to pathogenic and non-pathogenic mycobacterial infection and to determine whether it affects intracellular mycobacterial survival.

Firstly we sought to determine whether *OASL* upregulation was dependent on pathogenicity and virulence of mycobacteria by using the non-pathogenic *M. bovis* BCG strain, as well as the closely related hypo- and hypervirulent *M. tuberculosis* strains (Fig.1). Using THP-1 macrophages for our infection model [25], our results indicate that OASL is upregulated 24 h p.i in response to pathogenic *M.tb* infection only. Although *M. bovis* BCG does not induce its expression, it is likely that this occurs at a later time-point. The fact that we observe intracellular replication of *M. bovis* BCG 96 h after *OASL* silencing supports this (Fig. 2C). Our results are in accordance with other work indicating an upregulation of *OASL* in response to pathogenic mycobacteria (*M. leprae*) and not in response to non-pathogenic *M. bovis* BCG infection in THP-1 cells [13]. After phagocytosis, mycobacterial species mediate the phagosome breach, a process facilitated by the ESX-1 secretion system [26] which is only present in pathogenic mycobacteria. This leads to content leakage in the form of dsDNA into the host cytoplasm which was shown to trigger cytoplasmic DNA sensing receptors and induce type I IFN production [26, 27]. The secreted type I IFNs then act in an autocrine and paracrine fashion which induces OASL transcription through the STING/TBK1/IRF3 axis during the initial stages of infection [5]. *OASL* could therefore be an early response gene involved in host defence mechanisms against pathogenic mycobacteria.

The next step was to determine whether mycobacterial growth is affected by the absence of OASL, and in doing so evaluate the extent to which it contributes to host defence. We silenced *OASL* with two siRNAs targeting different sequences (denoted siRNA#1 and siRNA#2), and allowed 96 h for infection with each of the strains (Fig.2). We observed a similar trend whereby CFUs for all three strains increased significantly after knock-down 96 h p.i. Our results mimic those observed when OASL is inhibited during viral infection, resulting in enhanced viral replication and survival by reducing the type I interferon response [4].These results are however in contrast with work which observed that *M. leprae* survival decreased following *OASL* knock-down in the same cell type [13]. The authors suggest that during *M. leprae* infection, mycobacterial DNA activates cGAS, which leads to production of cGAMP, a STING agonist, which consequently activates OASL. OASL then inhibits cGAS which limits the type I IFN response and promotes intracellular M. *leprae* growth [13]. The reasons for this remain unclear since work associated with OASL signaling during bacterial infection is sparse, and to date only includes this study and the study on *M. leprae* [13]. To further complicate matters, upon entry of *M. tb* into the host cell, cyclic diadenylate monophosphate (c-di-AMP), a key mycobacterial pathogen-associated molecular pattern (PAMP), was found to associate with STING to drive type I IFN responses, and that c-di-AMP, not cytosolic DNA alone, is a ligand for IFN activation during *M. tb* infection [28]. There is no ortholog of diadenylate cyclase (and therefore no c-di-AMP) in the *M. leprae* genome, thus IFN signaling in both infection models differ, including how this affects OASL function.

Cytokines play a fundamental role in controlling mycobacterial infection without promoting uncontrolled and damaging inflammatory responses. Since OASL is induced following the onset of the inflammatory response, we investigated whether OASL knock-down affected the expression of various cytokines and chemokines as a possible explanation for the observed unrestricted intracellular growth (Fig. 3).

Although various cytokines and chemokines were assessed, our main findings show that both TNFα and IL-1β secretion is significantly reduced following OASL knock-down (Fig. 3A and B), which is in agreement with was observed elsewhere [4, 29]. Recently, both these cytokines were described as hallmarks of the innate immune response by macrophages that contribute to “trained immunity”, and are essential in controlling mycobacterial growth [30]. Finally, OASL knock-down reduced MCP-1 secretion, however only during infection with pathogenic mycobacteria. Evidence suggests that decreased MCP-1 secretion together with lowered TNFα contribute to the unrestricted growth and dissemination of mycobacteria [29]. OASL knock-down during M. leprae infection reflected a strikingly similar result [13]. OASL therefore appears to enhance pro-inflammatory cytokine and chemokine secretion during mycobacterial infection, however the mechanism behind this modulation is uncertain.

Although OASL is overexpressed following infection with *M. tb*, future studies should test whether a similar result is observed through a vector-based overexpression system. Additionally, it should be determined whether the induction and function of *OASL* is dependent on type I IFNs and whether the blockade of type I IFN signaling increases or decreases bacterial loads. Since IL-1β,TNF-α and MCP-1 show the greatest effect after OASL knock-down, neutralisation of each of them (together or alone) should affect *M.tb* intracellular replication to some degree. Although our study does not uncover the mechanism by which *OASL* enhances secretion of the pro-inflammatory mediators during mycobacterial infection, it does uncover for the first time that *OASL* supresses mycobacterial intracellular replication. OASL may no longer be associated exclusively with antiviral responses, but is now associated with antimicrobial responses and host defence mechanisms in the macrophage.

## Ethical Approval

Ethical approval was not required.

## Figure Legends

**Supplementary Figure S1**. A. Measurement of equal uptake of pathogenic and non-pathogenic mycobacteria in THP-1 cells 4h p.i. **B.** Confirmation of OASL knock down using two siRNA complexes with different targets as measured by qPCR in THP-1 cells. **C**. IL-10 secretion as measured through Luminex revealed no significant changes in IL-10 secretion after OASL silencing.

