## Supplementary figures and images for "2′-5′-Oligoadenylate synthetase-like protein inhibits intracellular *M. tuberculosis* replication and promotes proinflammatory cytokine secretion"

### Supplementary figure 1

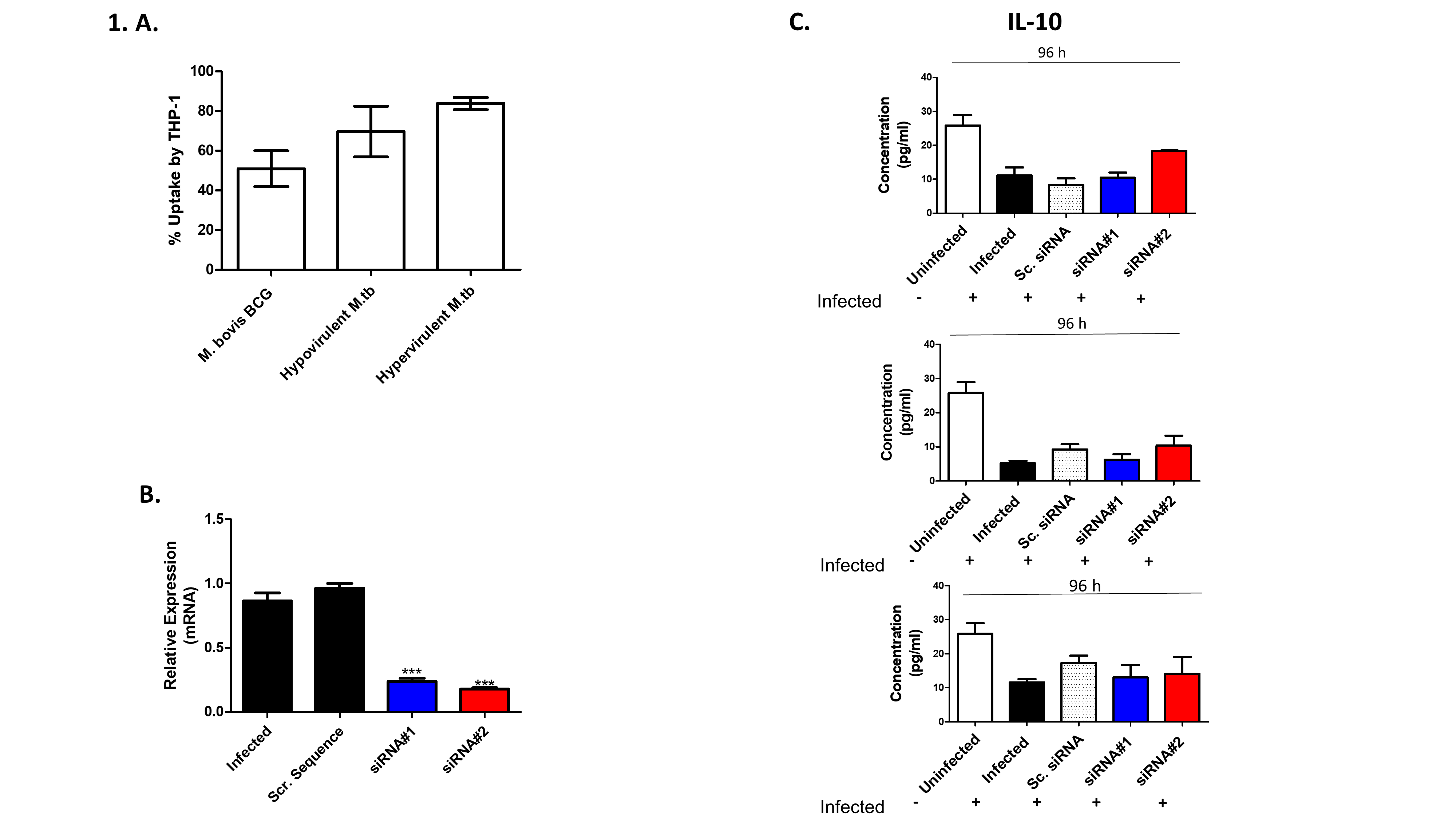
